# Altered Myocardial Structure in Post-natal Tetralogy of Fallot – A Substrate for Interpretation of Ventricular Function and Dysfunction?

**DOI:** 10.64898/2026.04.29.721688

**Authors:** Vaishnavi Sabarigirivasan, Joseph Brunet, Hector Dejea, Adrian Crucean, Anusha Jegatheeswaran, Giorgia Bosi, Theresa Urban, Lisa Chestnutt, Joanna Purzycka, Paul Tafforeau, Mark Friedberg, Peter D Lee, Andrew C Cook

## Abstract

**BACKGROUND:** In tetralogy of Fallot (ToF), changes in right ventricular function (as assessed by strain or TAPSE) reflect altered myocardial structure. Direct three-dimensional anatomical evidence supporting these changes remains limited. The objective of this study was to non-destructively characterize myocardial architecture in pediatric ToF hearts using Hierarchical Phase-Contrast Tomography (HiP-CT) and structure tensor analysis.

**METHODS:** Twenty ToF and control pediatric hearts were imaged at the European Synchrotron, ESRF. Myocyte orientation was assessed through structure tensor analysis and distributed high-performance computing. A region-specific framework was developed for analysis of the right ventricle. The predominant direction of myocardial aggregates (their helical angle) was compared across ventricular regions.

**RESULTS:** Significant differences in orientation were found in all ToF segments vs controls (left ventricle, right ventricular inlet, right ventricular outflow tract, septum; p < 0.001). Myocytes in the ToF right ventricular inlet were more circumferential overall, with regional heterogeneity. Contrary to traditional models, no discrete ‘middle layer’ was found in the ToF right ventricle; instead, a shift towards more circumferentially orientated myocytes and disrupted septal and outflow components was observed. Right ventricular contribution to the septum was greater in ToF (47.3% vs 34.0%; p = 0.0026), with extension of ventricular insertion points disrupting septal architecture. There were more longitudinally oriented myocytes in the ToF right ventricular outflow tract, consistent with hypertrophied septoparietal trabeculations. Left ventricular structure in ToF demonstrated a greater proportion of circumferentially oriented myocytes compared to controls.

**CONCLUSIONS:** We reveal profound alterations in ToF myocardial organization which broadly align with clinical observations and provide the first open-access HiP-CT congenital heart disease data as a basis for future computational modelling.

**Clinical Perspective:** Whole-heart HiP-CT demonstrates a loss of normal LV-RV distinction in the ToF myocardium, alongside extensive septal disarray. These findings provide a structural substrate for RV dysfunction, ventricular-ventricular interaction, and arrhythmogenesis in ToF, challenging traditional layer-based models of ventricular myocardium. Understanding myocardial organization as a continuous, developmentally patterned three-dimensional structure is essential for accurate interpretation of ventricular mechanics and disease progression. Although HiP-CT imaging is not applicable in vivo, the structural phenotypes identified in this study generate testable hypotheses for clinical imaging. Future work should focus on correlating ex-vivo measures with in-vivo imaging markers derived from cardiac magnetic resonance, including strain, TAPSE, and assessment of ventricular interactions. Investigating myocardial phenotype across the life-course, from fetal life to adulthood, paired with multi-omics mapped to these three-dimensional datasets, may help elucidate mechanisms underlying myocardial remodeling in ToF and support the development of novel therapeutic approaches.

## Introduction

In repaired Tetralogy of Fallot (ToF), despite excellent early survival, long-term morbidity and mortality continue to be important medical burdens^1,2^. Right (RV) and left (LV) ventricular function are important determinants of clinical outcomes in this population and contribute to the increasing burden of right heart failure in adults with congenital heart disease^3^. Outcomes such as exercise capacity and the progression of RV dysfunction vary widely between patients, and the mechanisms driving this variability are poorly understood. Currently, there is no useful anatomic data available to explain the variability in RV function and its clinical consequences, interpret the appearances seen on imaging, nor for computational modelling of this process.

Traditional approaches for investigating myocardial structure have relied on destructive methodologies, such as dissection by peeling, or histological analysis, which are not transferable to computational modelling^4,5^. While ex-vivo diffusion tensor imaging is a validated, non-destructive alternative, it is limited by spatial resolution and provides a surrogate rather than a direct measurement of myocyte orientation^6,7^. Hierarchical Phase-Contrast Tomography (HiP-CT) enables non-destructive imaging of intact post-mortem human organs at near-cellular resolution using the Extremely Brilliant Source at the European Synchrotron, ESRF^8^. Previous studies using synchrotron-based imaging techniques have visualized myocardial structure in whole fetal specimens^9–13^, but only HiP-CT has the potential to achieve this in organs of any size, including adult hearts^14^.

Our primary aim was to determine whether this new form of imaging, together with structure tensor analysis, can provide the necessary detail to explore myocardial architecture in archival pediatric specimens. In parallel, imaging and analysis workflows were optimized to minimize scan time, data size, and computational resources for analysis. A secondary aim was to produce fine-grained, open-access data of myocardial architecture in ToF for future computational modelling and comparison with clinical parameters.

## Materials and Methods

### Specimen Description

Five hundred and fifty ToF hearts from the UCL Cardiac Archive Biobank and Birmingham Children’s Hospital were screened, from which eleven were selected for imaging. Samples were chosen to represent a broad range of phenotypes, provided they were well preserved and within size limits (105 × 105 × 167 mm) to optimize imaging time and data volume. Nine size-matched structurally normal control hearts from the same collections were also included (ethical approval: REC 22/PR/0906). Specimen details are provided in Table 1.

**Table 1:**
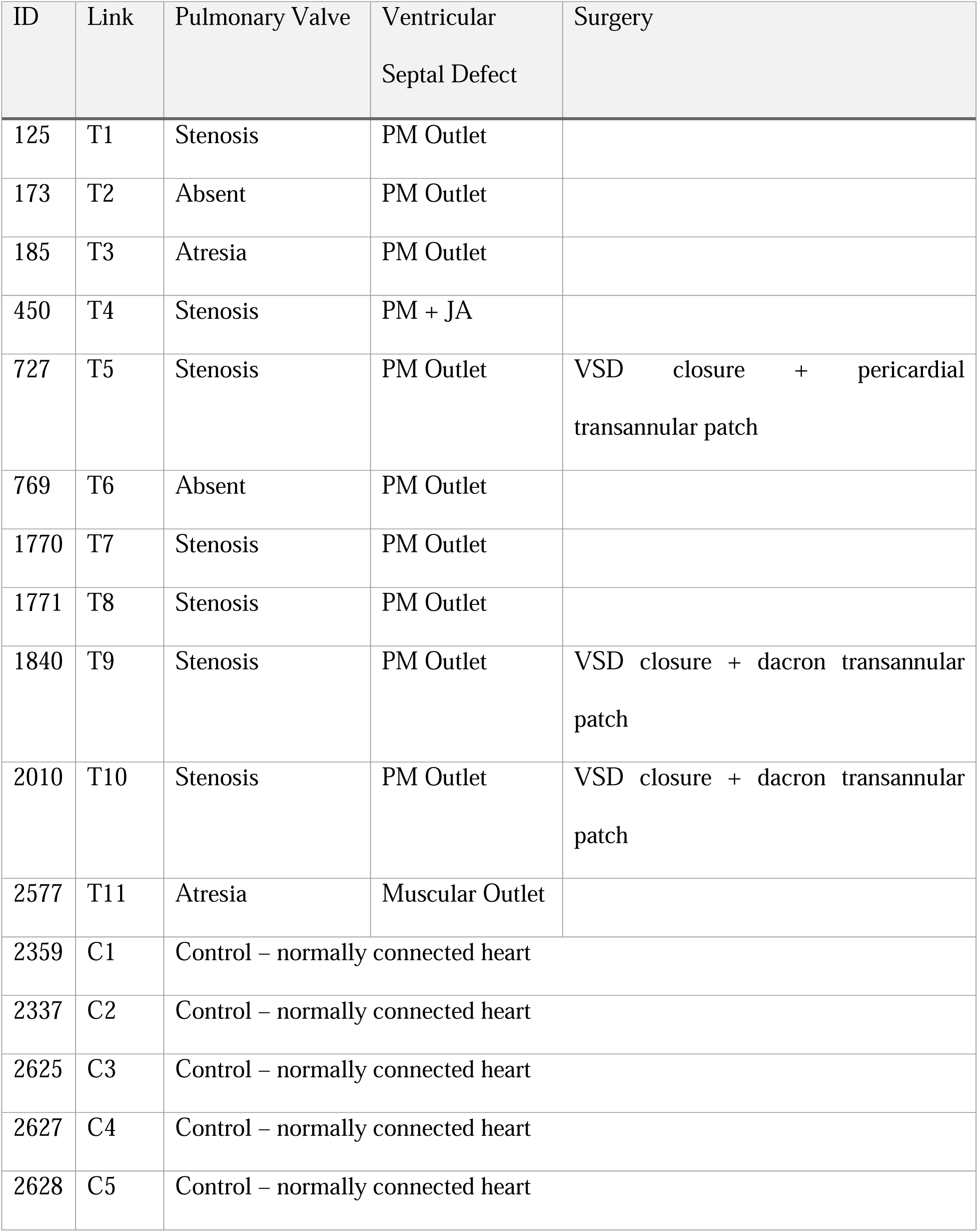
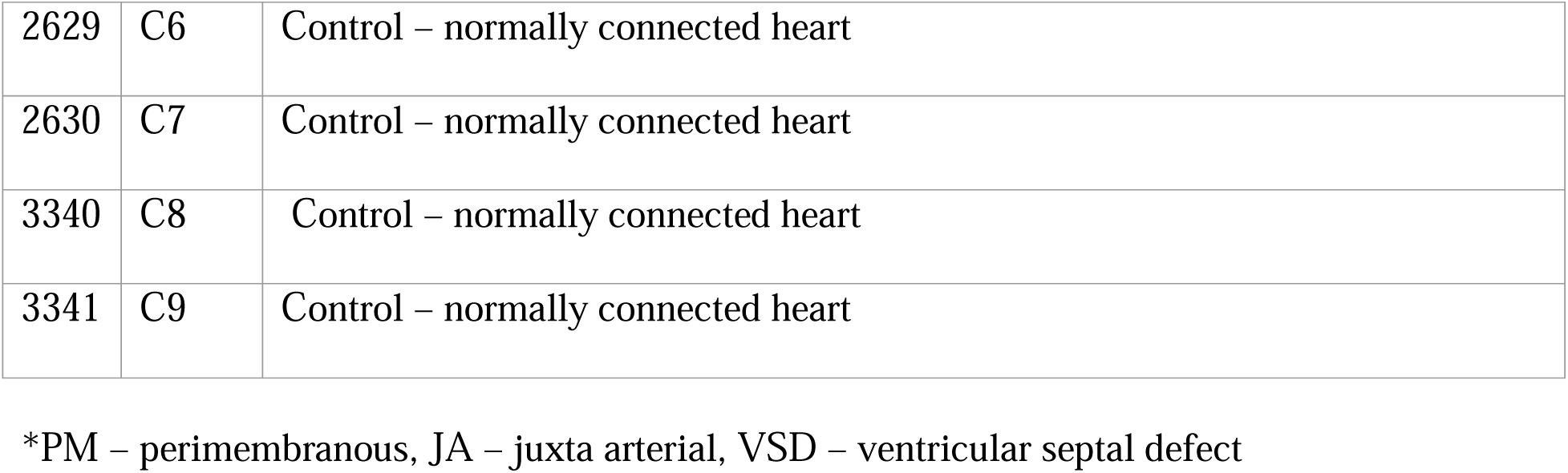
Morphologic characteristics of specimens included in this study.

All archival postmortem samples were assigned unique UCL Biobank IDs and transported to ESRF in compliance with the UK Human Tissue Act (Licence 12220), biobank standard operating procedures, and French Ministry of Higher Education and Research authorization (No. IE-2023-2966). In the United Kingdom, there has always been a requirement for consent for postmortem examination, including use for “research and education.” Many of these samples are now older than 60 to 80 years, and it is not considered appropriate for us to ask families for consent for publication in retrospect. Instead, families are allowed to withdraw consent for use of material at any time. The use of such material in the United Kingdom is regulated under the UK Human Tissue Act 2006 as “existing material”. We have confirmed that publication, including images if anonymized, is allowed under UK Human Tissue Authority legislation.

### Sample Preparation and Data acquisition

All hearts had been fixed and stored, long-term, in 10% formalin in standard archival fashion. Samples were stabilized and degassed in 4% formalin-agar and prepared for HiP-CT imaging at beamline BM18 (European Synchrotron, ESRF, Grenoble, France)^15^ according to the protocol of Walsh et al.^8^ and Brunet et al.^16^ (modified for imaging in 4% formalin). Whole-organ scans were performed at voxel sizes ranging from 7-20μm depending on specimen size. Full details can be found in supplementary materials.

### Contrast-to-Noise Ratio Quantification and Image Quality

Contrast-to-noise ratio (CNR) was quantified on the full-field images using ImageJ v1.54g^17^ using the following formula:

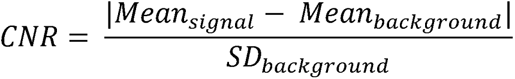

CNR was measured in the ventricular myocardium in five randomly selected areas in each dataset. The formalin-agar gel was taken as the background.

### Ventricular Wall Thickness

Ventricular free wall thickness was quantified using a custom Python script (see Code Availability). In five evenly spaced slices, transmural lines were drawn perpendicular to the endocardial surface. Thickness values were interpolated between slices to estimate wall thickness across the full volume, and the mean thickness was calculated for each dataset.

### Cardiomyocyte aggregates orientation

Myocardial organization was assessed using the helical angle (HA), which describes the predominant direction of small aggregates of (approximately 2-5) cardiomyocytes, relative to the circumferential plane of the ventricular wall. A HA of 0° indicates a circumferential orientation, while positive and negative values indicate longitudinal orientations in opposing directions. Cardiomyocyte orientation was quantified and visualized with a gradient structure tensor approach using in-house, open-source code, Cardiotensor^18^ (see Code Availability). Full details of the structure tensor method can be found in supplementary materials.

### Implementing a dual co-ordinate system for analysis of the Right Ventricle

In previous diffusion tensor MRI (DT-MRI) and phase-contrast X-ray studies the ventricular myocardium has been analyzed almost exclusively using a long-axis centreline within the LV, which may bias mapping of regions outside or distant from the LV axis. To illustrate this effect, mapping was performed from three different centrelines within the heart and compared qualitatively in control and ToF hearts (Figure 1). Having confirmed our suspicion of potential bias, a local coordinate system was defined for each region of interest: RV inlet, RV outlet and combined LV inlet/outlet. The LV axis was established between the center of the mitral valve and the LV apex, as previously defined in the literature for synchrotron imaging and diffusion tensor MRI studies^10,11,13,19^. RV inlet and outlet regions were divided by a line drawn from the anteroseptal commissure to the RV apex, with axes established between the center of the tricuspid valve or pulmonary valve and the RV apex respectively (Figure 2). Segmentation of structure tensor data was performed in 3D using VGStudioMax 2024.4 software.

**Figure 1:**
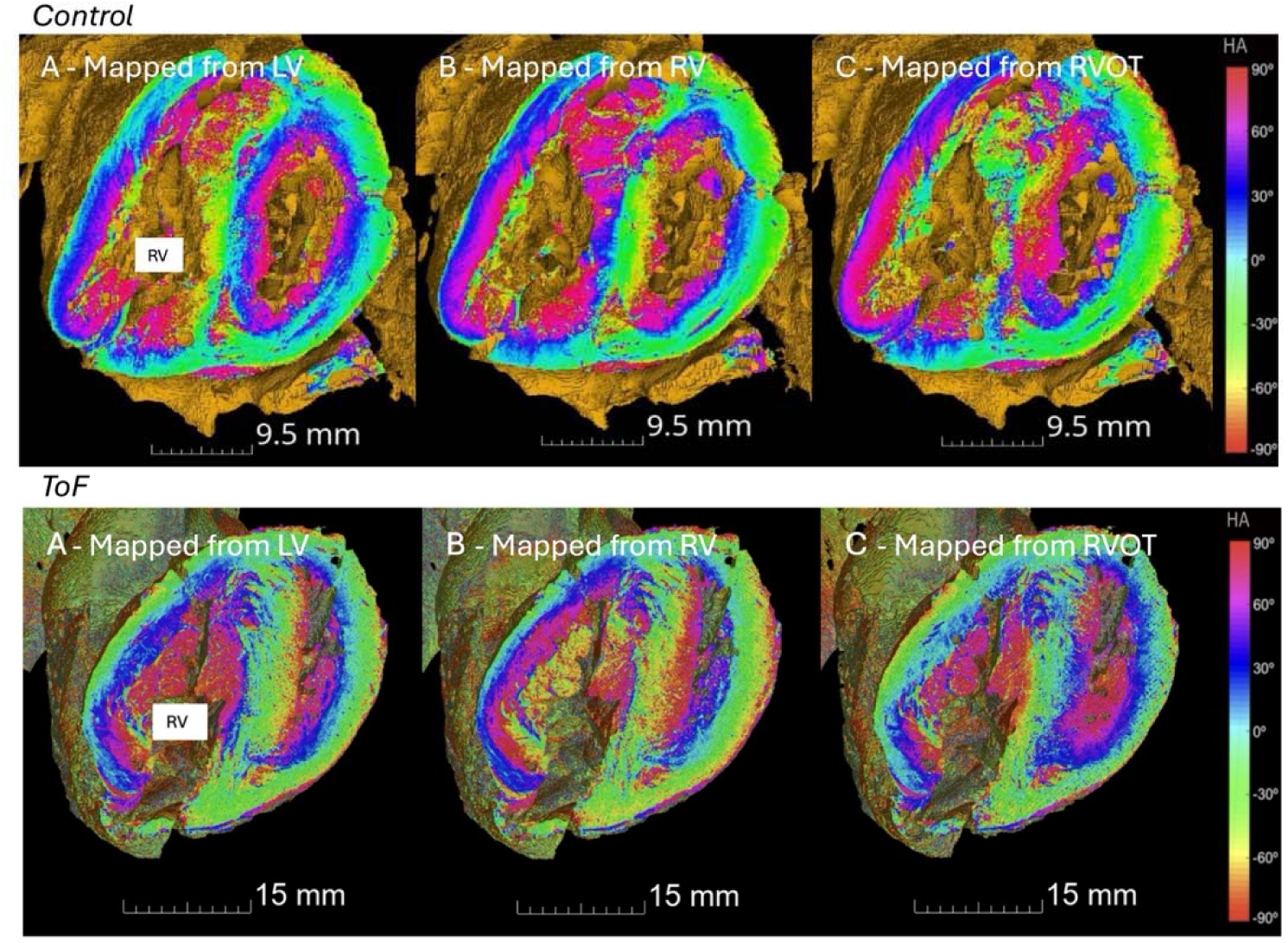
Comparison of structure tensor analysis of helical angle (HA) in mid-ventricular slices using different mapping axes in a control and ToF sample. A - LV axis, B - RV inlet axis, C - RVOT axis.

**Figure 2:**
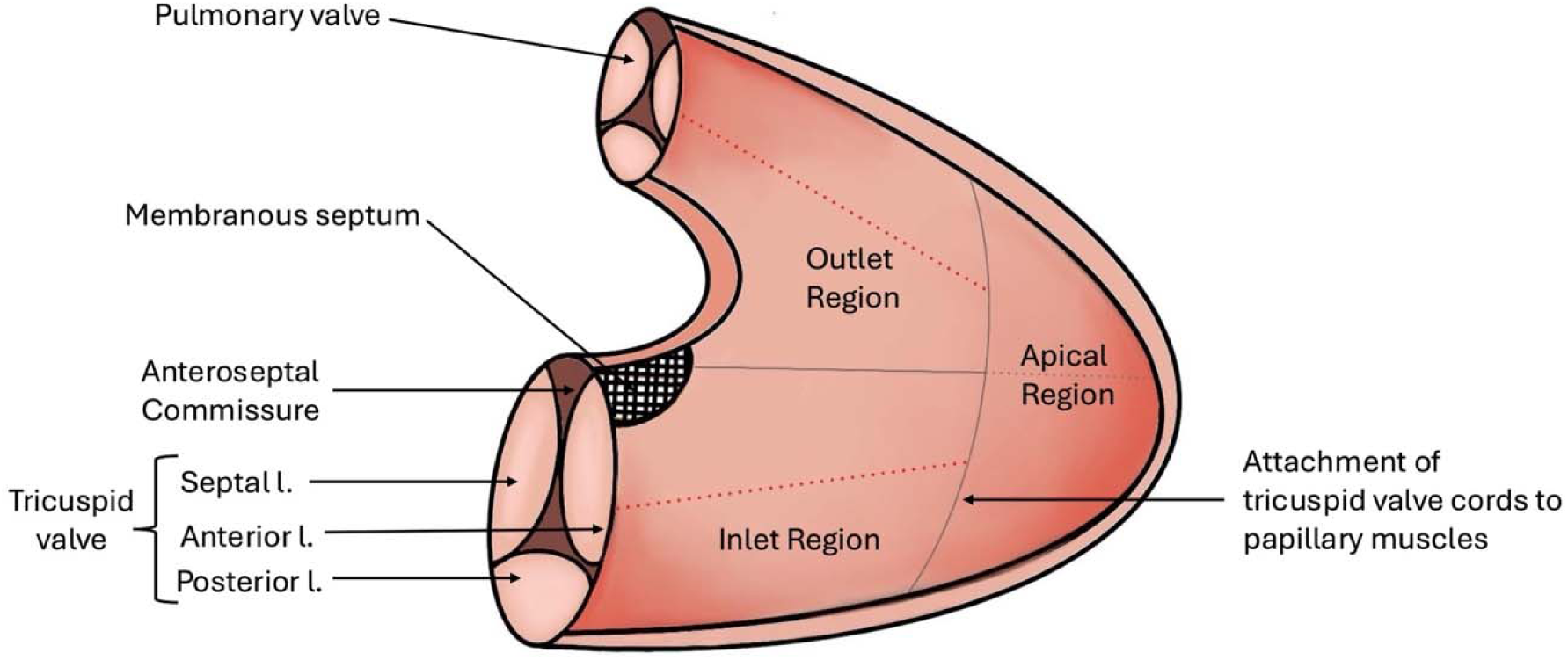
Proposed method of right ventricle (RV) analysis. Using one line extending to the RV apex from the anteroseptal commissure, and another at the attachment point of the tricuspid valve cords to the papillary muscles, the RV is divided into inlet, outlet, and apical regions. Dashed lines - centers of the local coordinate systems in the inlet and outlet regions extending to the RV apex from either the tricuspid valve or the pulmonary valve, respectively.

### Quantification of Left and Right Ventricular Contribution to the Interventricular Septum

To assess the relative contribution of the LV and RV to the interventricular septum, a combined anatomical and geometric approach was employed, adapted from Zhang et al^20^. High-resolution HiP-CT datasets enabled clear visualization of septal regions in continuity with ventricular free walls based on myocyte orientation, with the region of circumferentially orientated aggregates taken as the transition point between ventricles (Figure 3).

**Figure 3:**
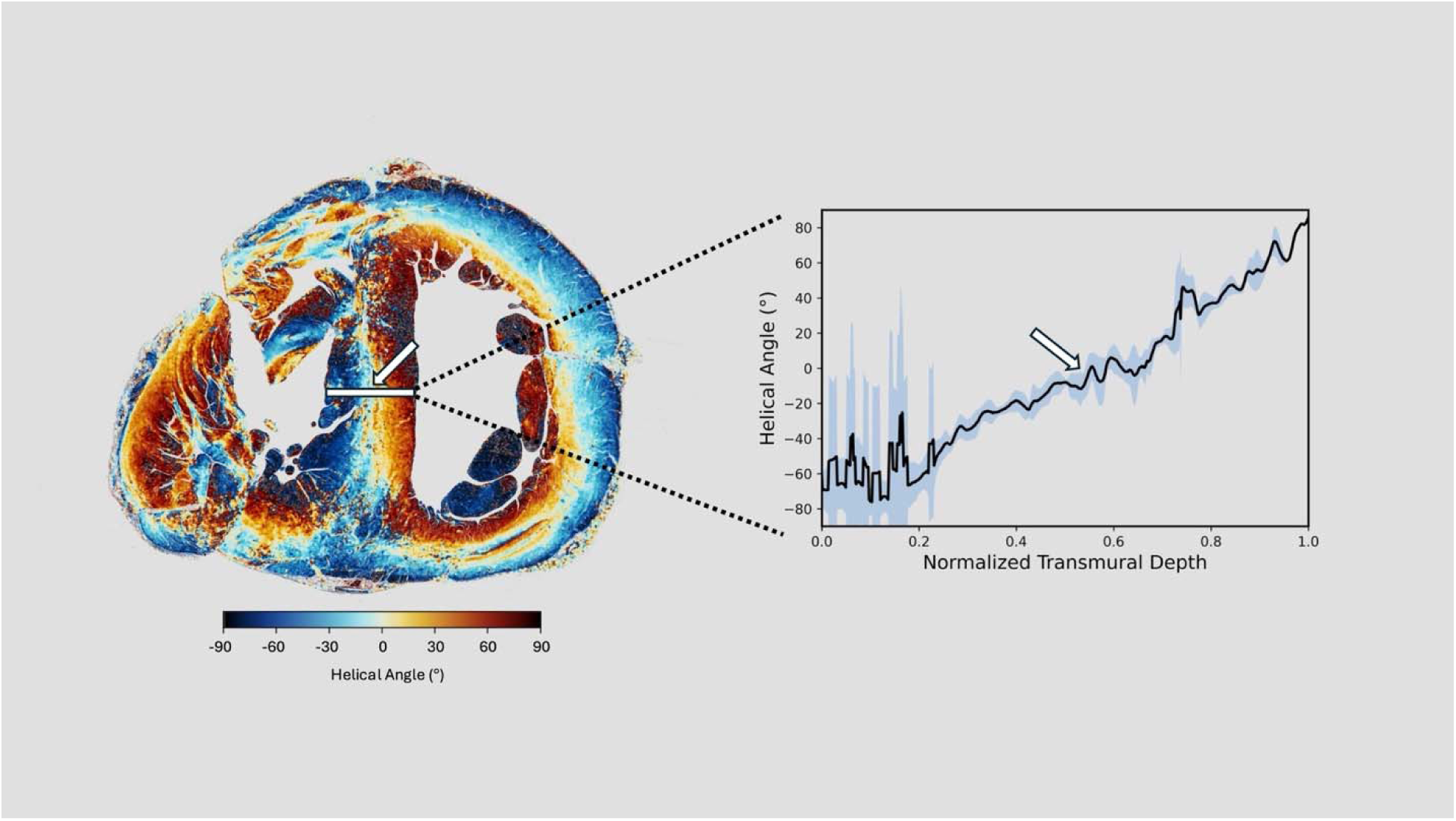
Helical angle change across the interventricular septum in a ToF heart demonstrating equal contribution of right and left ventricles. The graph demonstrates the change in helical angle across the septum. Red arrows point to the region of circumferentially orientated cardiomyocytes (HA = 0°) - The transition point between the left and right ventricles.

### Colourmap selection for Helical Angle Visualization

Helical angle visualization was facilitated using a cyclic, perceptually uniform colormap, in line with Society for Cardiovascular Magnetic Resonance recommendations^21,22^ (details in Supplementary Materials).

### Statistical Analysis

Statistical comparisons were performed using t-tests and one-way ANOVA with appropriate post-hoc testing; full details are provided in Supplementary Materials.

## Results

### Contrast-to-Noise Ratio

CNR measurements are summarized in Supplementary Table S3. Notably, samples previously exposed to Fomblin for DTMRI imaging showed significantly lower CNR than unexposed samples (5.4 ± 1.2 vs 9.4 ± 3.7; p = 0.002), despite appearing entirely normal on gross inspection.

### Ventricular Free Wall Thickness

Ventricular free wall thickness, indexed to ventricular length, was compared between ToF and Control groups. A significant difference in wall thickness between the LV and RV was observed in the Control group (p = 0.015), however in ToF, this difference was lost (p = 0.174) (Figure 4).

**Figure 4:**
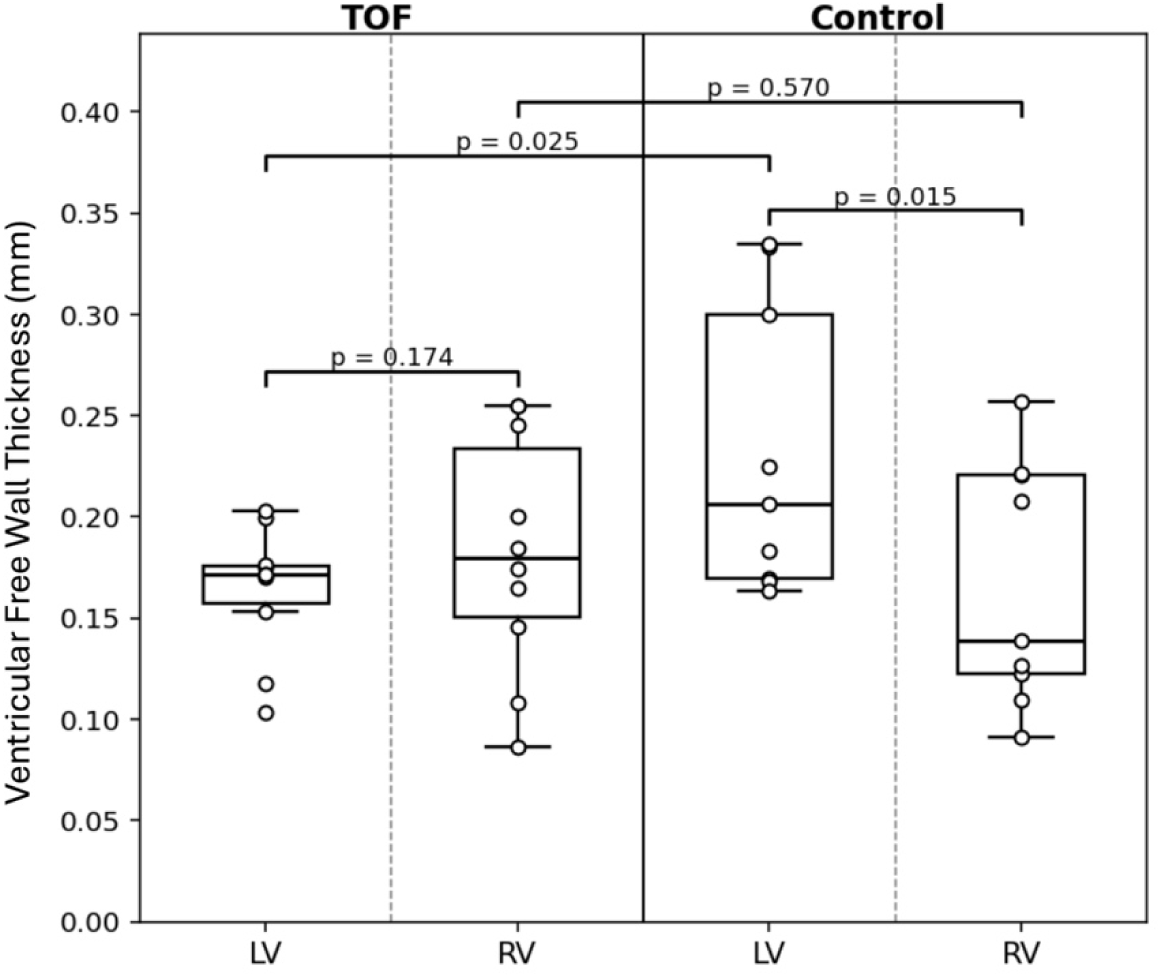
Box and whiskers plots show the thickness of the left and right ventricular free walls in control and ToF hearts in datasets indexed against ventricular length.

### Myocardial Architecture

Myocardial organization could be clearly visualized in the whole heart volume, and an underlying, smooth linear transition from endocardium to epicardium was qualitatively demonstrated in both control and ToF hearts in both ventricles (Figure 5). One-way ANOVA to assess regional differences in myocyte orientation between control and ToF groups showed statistically significant differences across all regions: LV (F = 472.2, p <0.001), septum (F = 546.9, p <0.001), RV inlet (F = 1884.7, p <0.001), and RVOT (F = 267.9, p <0.001), shown in Figure 6.

**Figure 5:**
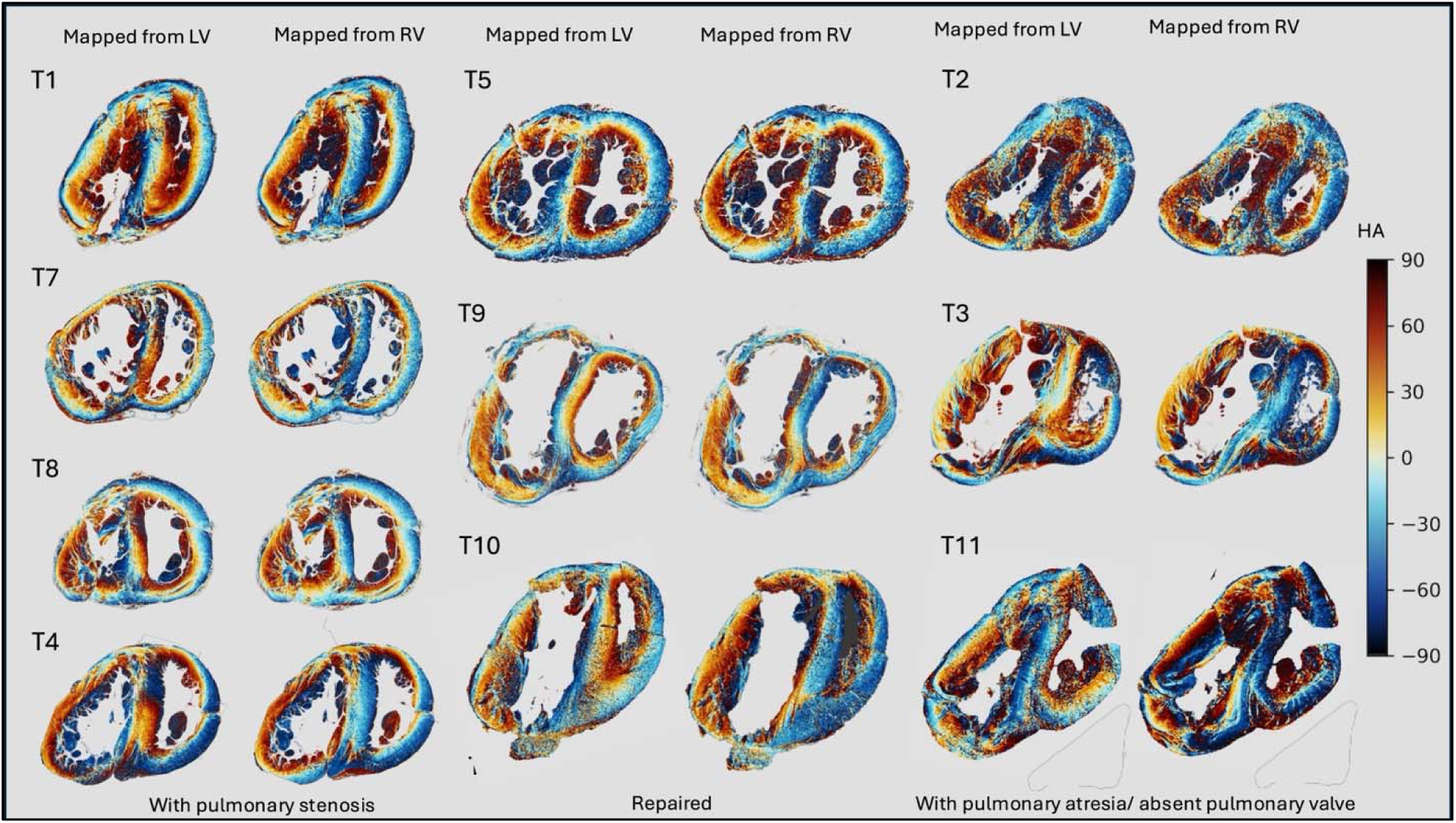
Mid-ventricular slices of ToF specimens mapped from the left ventricle (LV) and the right ventricular (RV) inlet, grouped by outflow tract morphology.

**Figure 6:**
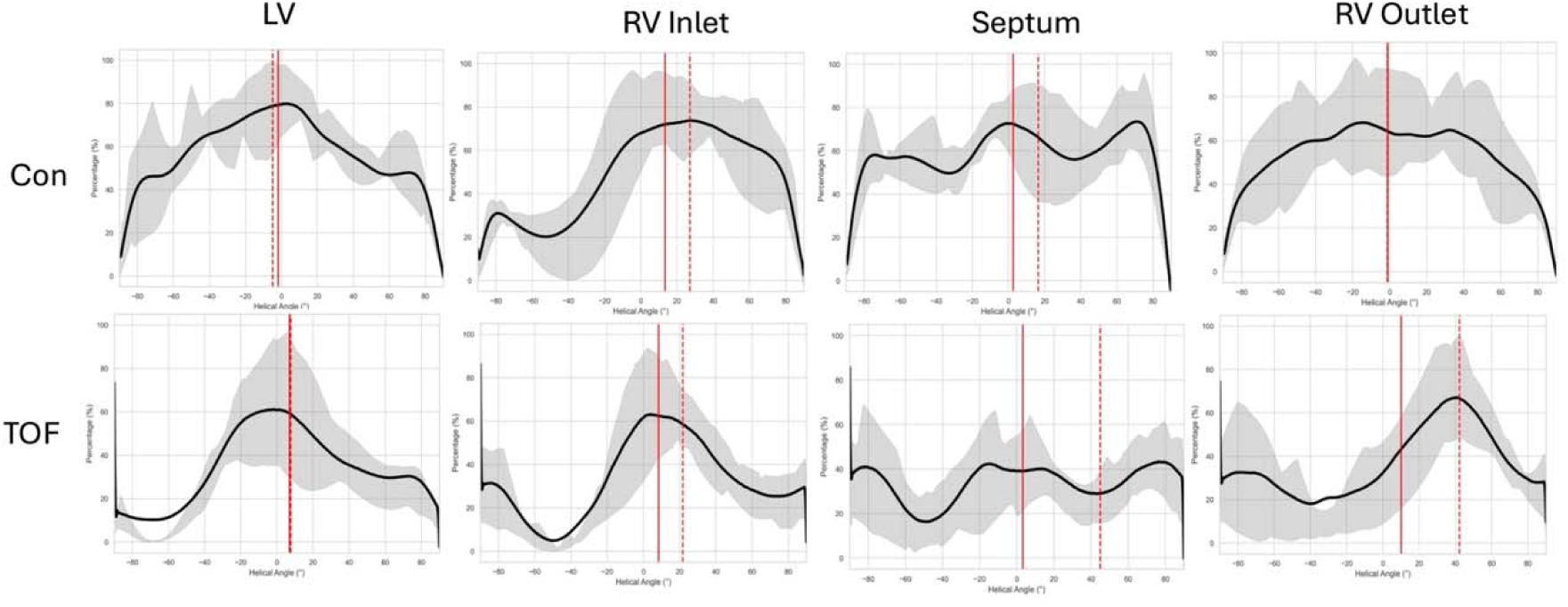
Helical angle frequencies displayed as histograms in different regions of the heart and in control versus ToF samples. Interquartile range is shown as the shaded region. Solid red lines mark the numerical means, while circular means are marked by the dashed line.

#### Right ventricular structure in ToF

In control specimens, RVs were heterogeneous in structure both between samples and transmurally in the same sample when compared to the LV. Despite regional heterogeneity, there was an underlying smooth transition in predominant cardiomyocyte orientation from the endo- to epicardium. A significant difference was observed in the distribution of HA between LV and RV inlets (mean difference = –0.0963, *p* < 0.001), indicating distinct structural profiles.

In ToF specimens, the RV demonstrated greater inter-specimen variability compared to controls, again with regional heterogeneity. There was no statistically significant difference in ToF between the HA distributions of the LV and RV inlet (mean difference = −0.0095, p = 0.088).

In a subset of ToF hearts, the RV closely resembled the LV, exhibiting smooth, concentric transmural HA progression from endocardium to epicardium (e.g. Figure 5, T1). Other specimens displayed a markedly heterogeneous pattern of alternating stripes of aligned myocytes, producing a banded appearance (e.g. Figure 5, T3).

#### The interventricular septum

In the short axis of the normal heart, the insertion points of the LV and RV into the interventricular septum were well-defined triangles, continuous with the surrounding alignment of myocytes of each ventricle. There was clear continuity of myocyte aggregates between the ventricles along their diaphragmatic border, below the inferior insertion point.

In contrast, ToF specimens demonstrated marked variability in the insertion of the ventricles into the interventricular septum, both in the degree of disarray, and in the extent to which the insertion points extended medially into the septal myocardium in short axis (Figure 7). In most ToF hearts, the inferior insertion disrupted the normal inferior continuity of myocyte aggregates between ventricles, instead forming an elongated triangular region, often extending from the RV. The inferior insertion points in all specimens extended into the septum, occupying up to one-third of the total length of the septum in some (T2,T3,T4,T11). Superior insertion points either showed similar disarray and extension into the septum (T1,T3,T4,T7,T11), or a more normal pattern with a discrete region of localized disarray (T2, T5, T8, T9, T10). When displaying the interventricular septum en-face, abnormal insertion points were seen to extend into the septum from base to apex, throughout the length of the RV (Supplementary Figure S2).

**Figure 7:**
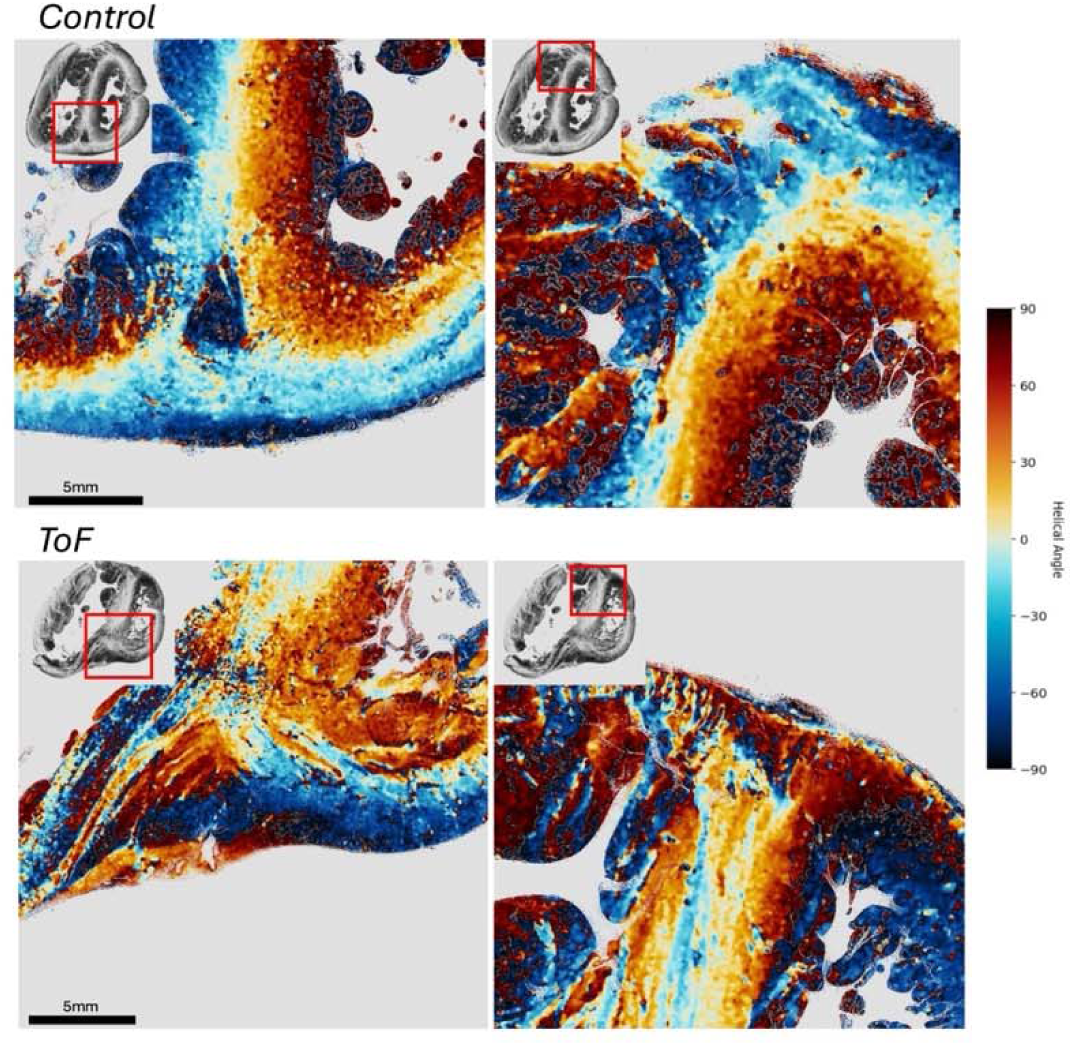
Insertion points of ventricles into the interventricular septum in a control and ToF sample depicting the increased extension of the insertion points into the septum in ToF samples.

Operated specimens (T5,T9,T10) exhibited the most control-like septal morphology. Whilst disarray was present in the insertion points, their extension into the septal myocardium was minimal and the inferior continuity of myocyte aggregates was largely preserved.

Finally, in ToF specimens, there was an increased contribution of the RV to the mid-septal wall compared to controls (47.3% ± 5.9% versus 34.0% ± 9.4%, p = 0.0026).

#### The Right Ventricular Outflow Tract (RVOT)

The ToF RVOT demonstrated a higher percentage of longitudinally orientated myocytes compared to control specimens, in which the HA distribution showed predominantly circumferential orientation. All inter-phenotype comparisons in the RVOT reached statistical significance, with ToF-PA demonstrating almost exclusively longitudinally orientated myocytes (Supplementary Figure S3).

#### The Left Ventricle

The LV of control specimens demonstrated a broad HA distribution, and a peak of circumferentially orientated myocytes at 0°, consistent with a gradual change in cardiomyocyte orientation across the wall. The ToF LV also demonstrated a peak of circumferentially orientated myocyte aggregates, with a peak close to zero, but within a narrower distribution of helical angles (Figure 6).

## Discussion

Long-term morbidity remains a significant burden in ToF, particularly due to RV dysfunction. While hemodynamic factors such as chronic pulmonary regurgitation and myocardial pathology such as fibrosis are recognized as risk factors for RV dysfunction^3^, they do not adequately explain or predict clinical course. This knowledge gap complicates risk prediction, management, and counseling. Previous studies using computational modelling demonstrate that reduced RV contractility is a strong determinant of exercise capacity, more than the degree of pulmonary regurgitation^3^. The myocardial structure of the RV, and its changes in ToF, may be a significant determinant of RV contractility, overall performance, and response to adverse hemodynamic load, yet these are poorly understood, and the tools to adequately quantify these in the intact heart post-mortem have been limited.

### HiP-CT Imaging in the Assessment of Myocardial Structure

HiP-CT enables exceptional resolution preserving whole-organ 3D spatial context, effectively providing 3D whole heart histology and allowing comparison of samples from fetal life to adulthood. In contrast, conventional examination of myocardial structure in pediatric hearts has been challenging, requiring destructive dissection or ‘peeling’, or else serial sectioning and analysis based on limited and potentially misaligned 2D data^4–6^. Both risk anatomical misinterpretation and lack the 3D context required for modelling disease in-silico.

### An Anatomically-Informed Approach to Assess Myocardial Structure in the Right Ventricle

This study introduces two novel concepts for the analysis of myocardial architecture: a dual co-ordinate system for RV analysis and the use of whole segmented volumes (RV, LV and ventricular septum). The American Heart Association’s 17-segment LV model has improved consistency and correlation of anatomy with clinical imaging^23^. The complex geometry and contractile mechanics of the RV have limited the development of similar standardized models. Existing RV segmentation schemes are themselves complex and lack grounding in anatomy and embryological development^24^. We propose a simplified, anatomically-informed, dual-axis model for RV analysis comprising inlet and outlet^25^. Separating this into an additional apical component was feasible, but was not the focus of the current study. Traditionally, the RV myocardium has been ‘mapped’ or interpreted using the cylindrical axis of the LV^7,19^, however, we demonstrate that this technique produces potential bias and creates different results when compared to mapping from either the RV inlet or RV outlet. Moreover, our volumetric approach to myocardial analysis takes advantage of the full depth of data available via HiP-CT, rather than focusing on representative slices at basal, mid-ventricular and apical levels, as reported previously.^11^

### The Right Ventricle

Significant debate surrounding mural architecture in the normal heart has led to the generally accepted view of discrete layers, three in the LV and two in the RV, or, for others, to the existence of a unique myocardial band^26,27^. Further experiments using pneumatic distention, circular knives and 2D histology have suggested that the structure of the LV is better described as a continuum^28,29^. More recently, DT- MRI, has been used to explore myocardial structure, and the presence and contribution of helical, intruding and ‘sheetlet’ angles is still under debate^7^.

Our whole-volume 3D analysis did not identify distinct layered boundaries in either control or ToF hearts; both groups demonstrated a gradual change in myocyte orientation from epicardium to endocardium, with comparable smoothness of transition. Control hearts showed statistically significant differences in HA distributions and wall thickness between the RV and LV, reflecting distinct physiological roles early in post-natal life. In ToF, these inter-ventricular differences were lost: the RV and LV exhibited no statistically significant difference in wall thickness or HA. This convergence likely reflects RV adaptation to altered haemodynamic load in ToF, generally accepted as a sequence of pressure overload, leading to sarcomeric replication in parallel, and RV thickening to normalize myocardial wall stress. An additional contributing mechanism may be interruption of the normal postnatal regression of RV mass, typically occuring within the first months of life with falling pulmonary vascular resistance, but may fail to occur in ToF^25,30^.

### The Interventricular Septum and Ventricle-ventricle Interactions

The normal postnatal emergence of LV dominance in the interventricular septum was absent in ToF. Instead, our data showed nearly equal LV–RV contribution, resembling fetal configuration at a time when ventricles experience comparable afterload^20^. Across all structures examined, the ToF septum displayed the greatest changes, particularly in the region of the superior and inferior insertion points of the ventricles into the septal myocardium, which occupied up to half the length of the septum in some cases. Similar regions are recognized as areas of late gadolinium enhancement in MRI studies, in diseases such as pulmonary arterial hypertension and hypertrophic cardiomyopathy and are thought to represent fibrosis^31,32^.

Septal architecture is critical for ventricular–ventricular interaction, and perturbations can propagate failure across chambers^33–35^. Our findings support this framework: the fetal-like septal configuration and extensive insertion points in ToF could impair the mechanical coupling required for normal LV function, further complicated by the presence of a postoperative VSD patch. This aligns with clinical observations reporting early postoperative electromechanical dyssynchrony and reduced RV longitudinal strain in infants with repaired ToF, with secondary impacts on LV performance through septal interaction^35,36^. If fibrosis does occur at the septal insertion points in ToF, this could further impair septal function and right-left ventricular interactions^37,38^. LV abnormalities in ToF are not isolated phenomena, but could be manifestations of ventricular interdependence driven by a developmentally abnormal, structurally disorganized septum. The microstructural disturbances demonstrated here provide an anatomic substrate for clinically observed altered biventricular mechanics. In control hearts overall, the LV exhibited a well-organized transmural HA gradient, with smooth transition from subendocardial to subepicardial aggregates. In ToF, the LV demonstrated more circumferentially orientated myocytes, in contrast to the normal LV, suggesting abnormal remodeling or intrinsic abnormalities. These findings hold substantial significance as LV dysfunction is an important risk factor for morbidity and mortality in repaired ToF, but the underlying mechanisms have been poorly understood^39^.

### The Right Ventricular Outflow Tract

In contrast to previous studies, we demonstrate that the ToF RVOT shows a shift towards longitudinal myocyte alignment compared to control hearts^4^. Significant differences in HA distributions in the RVOT were observed between ToF phenotypes. ToF-PA demonstrated the most longitudinally oriented HA profile, followed by ToF-PS, while APV hearts showed a greater circumferential component. This suggests that underlying hemodynamic burden may influence remodeling and disease progression in the ToF RVOT and is entirely consistent with the presence of abnormal septo-parietal trabeculations which are small and longitudinally aligned in the normal heart but become hypertrophied in ToF. Whether these structural changes contribute to overall RVOT dysfunction and the incidence of ventricular arrythmia, alongside surgical residua and anatomic isthmuses merits further investigation.

### Limitations

Although this study represents the largest high-resolution 3D characterization of the ToF myocardium to date, the sample size remains limited, particularly for phenotype-specific analyses (e.g., outflow tract variants), which may therefore be underpowered and should be interpreted with caution. Future work will require scaling through high-throughput synchrotron imaging pipelines capable of standardized, rapid acquisition across large cohorts. Advances in automated pre-processing, segmentation, and structure tensor analysis will be essential to manage computational demands and enable the generation of multicentre, high-resolution atlases of myocardial architecture in disease. Finally, some control hearts had previously been immersed in Fomblin for post-mortem diffusion tensor MRI. Although grossly normal, residual Fomblin was visible on HiP-CT and complicated myocardial analysis. We therefore do not recommend Fomblin use prior to HiP-CT.

## Conclusion

Abnormal myocardial architecture in ToF reflects both developmental and physiological influences. Our findings challenge the traditional ‘three-layer’ model and show a more heterogeneous RV inlet with ‘LV-like’ features, likely due to reduced postnatal regression and compensatory hypertrophy. The RVOT differs from the inlet, consistent with its distinct secondary heart field origin. The septum is also abnormal, with balanced RV–LV contribution and disruption extending from ventricular insertion points. Together, these results support a developmentally informed framework for studying ToF and its progression into adulthood.

## Code Availability

Ventricular wall thickness analysis: https://github.com/VaishnaviSabari/Ventricle-wall-thickness.

Cardiotensor package: https://github.com/JosephBrunet/cardiotensor.

## Data Availability

All reconstructed volumes are openly accessible through the Human Organ Atlas repository (https://human-organ-atlas.fr). Dataset DOIs corresponding to each donor are listed in supplementary table 1.

## Acknowledgements

We gratefully acknowledge ESRF beamtimes md1290 and md1389 on BM18, led by PD Lee, P Tafforeau, CL Walsh, et al., as sources of the data and for hosting the Human Organ Atlas Hub. We sincerely thank David Stansby and Guillaume Geisne for their support with data management and data upload, as well as Joanna Purzycka and Jean-Pascal Hograindleur for sample preparation and mounting at ESRF.

## Supplemental Material

Supplemental Methods

Table S1 – including datasets

Figures S1-S3

## Non-standard Abbreviations and Acronyms

APV: Absent Pulmonary Valve
CNR: Contrast-to-Noise Ratio
ESRF: European Synchrotron, Grenoble, France
HA: Helical Angle
HiP-CT: Hierarchical Phase Contrast Tomography
LV: Left Ventricle
PA: Pulmonary Atresia
RV: Right Ventricle
RVOT: Right Ventricular Outflow Tract
ToF: Tetralogy of Fallot

---

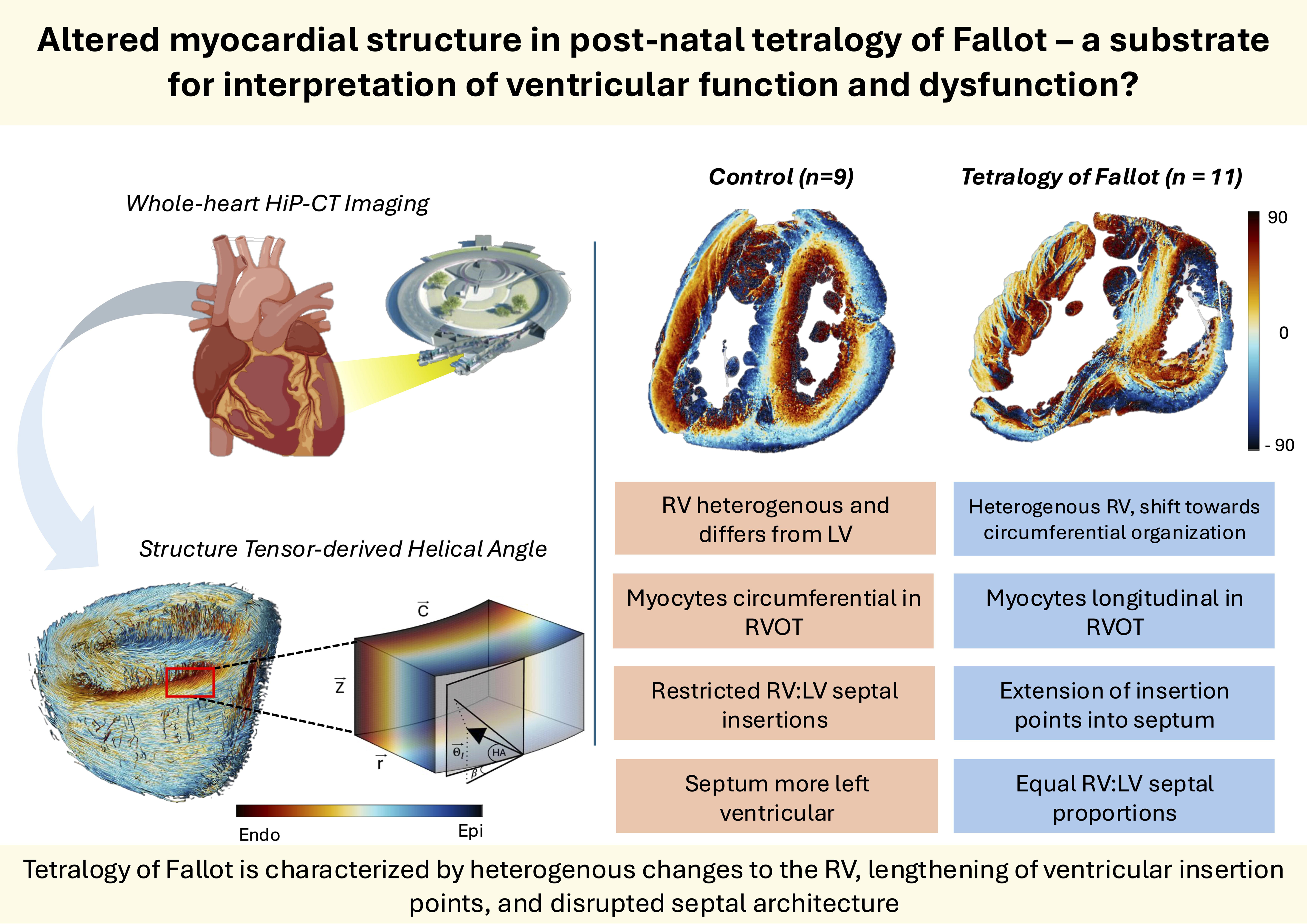

