## Supplementary for "Altered Myocardial Structure in Post-natal Tetralogy of Fallot – A Substrate for Interpretation of Ventricular Function and Dysfunction?"

### Supplementary Methods

#### Sample Preparation and Data Acquisition

Each heart was mounted in a container filled with 4% neutral buffered formalin and agar to stabilize the sample and prevent motion during acquisition. To minimize the formation of bubbles caused by radiation exposure, specimens underwent in-line degassing prior to imaging. The containers were then hermetically sealed until scanning started.

A filtered parallel polychromatic X-ray beam was used with a 178m source-to-sample distance and propagation distances of 10m (overview) and 1m/1.6m (local zoom)^1^. X-rays were converted to visible light using LuAG:Ce scintillators (Crytur, Czechia) of 2000 μm (overview) or 50 μm with a reflective layer or 100 μm (zoom) thickness, demagnified by Dzoom optic (overview) or magnified by a fixed x2 optic (zooms), and detected by an Iris 15 camera (Teledyne Photometrics, United States). Specimens were scanned entirely at voxel sizes of 19.19 μm (ToF with lungs), 10 μm (ToF without lungs, and ZCR normal specimens), and 7.032 μm (BCH normal controls), followed by local ‘zoom’ scans in regions of interest at 2.2 μm voxel resolution in all ToF samples and three control samples. The difference in overview pixel sizes is related to the size of the specimen. For each acquisition, a scan of a sealed container filled with 4% agar/formalin with the same parameters to serve as a beam reference for flat-field correction and correct low-frequency artefacts. Full scan parameters are in supplementary table S1.

#### Data processing and analysis

Data reconstruction was performed using tomography processing software (night-rail^2^, Nabu^3^) developed at ESRF, following the HiP-CT processing protocol as described by Brunet et al^1^. The night-rail framework integrates normalization, reference pairing, vertical concatenation (or helical). Ring artefact correction was applied on the reconstructed slices using an updated version of the Lyckegaard algorithm^4^.

#### Cardiomyocyte aggregates orientation

The structure tensor method is an approach for determining the orientation of imaged structures. It captures local orientation information around a point in space and is implemented using gradient calculations and neighborhood integration^5^.

In 3D, the structure tensor is represented as a 3x3 matrix. For a given volume, V, the structure tensor S at a point is computed as:

S = $\sum\nabla V{(\nabla V)}^{T}$,

Where $\nabla V={{[V}_{x}V_{y}V_{z}]}^{T}$ is the gradient of V, and the summation is performed over a specific neighbourhood around the point.

For gradient computation and integration, gaussian kernels are employed:

S = $K_{\rho}({\nabla V}_{\sigma}\left( {\nabla V}_{\sigma} \right)^{T})$,

Where $\sigma$ is the standard deviation of the Gaussian derivative kernel used for computing the gradient ${\nabla V}_{\sigma}$, and $\rho$ is the standard deviation of the Gaussian kernel $K_{\rho}$ used for integration. Here, $\sigma$ is referred to as the noise scale, which suppresses high frequency noise, while $\rho$ is the integration scale, chosen based on the size of the structures of interest^6^. At each voxel, this results in a symmetric 3x3 matrix:

$$S= \left[ \begin{matrix} S_{xx} & S_{xy} & S_{xz} \\ S_{xy} & S_{yy} & S_{yz} \\ S_{xz} & S_{yz} & S_{zz} \end{matrix} \right]$$

The structure tensor $\hat{T}$ is then decomposed into its eigenvectors and eigenvalues through the equation:

$$\hat{T}v= \lambda v$$

The eigenvectors $V_{i}$ (where $i \in\left\{ 1,2,3 \right\}$) represent orthogonal direction, with the smallest eigenvalue corresponding to the eigenvector pointing in the direction of lowest image intensity variation, assumed to be the predominant direction of the myocardial aggregate.

For this study, the orientation of the ventricular myocyte aggregates was assessed primarily through the helical angle (HA). The coordinate system was changed from a cartesian coordinate system to a cylindrical coordinate system by applying a 2D rotation to the x and y components of the vectors. The HA was calculated between the tertiary eigenvector and the local circumferential plane. Myocyte aggregates were considered circumferential if |HA| $\leq$ 20°, longitudinally positive if HA > 20°, and longitudinally negative if HA < 20°. Transmural profiles, from endo-to epicardium of HA were obtained in each region of interest and a linear regression fitting, y = β _1_ ·x + β _0_ , was applied to transmural HA profiles to characterize their linearity^5^.

Developing parameters for $\sigma$ and $\rho$ is essential to capture myocyte orientations without over-smoothing or amplifying noise^6^. Parameters for most samples was set to $\sigma$ = 1, $\rho$ = 5 at an approximate voxel resolution of 10μm, and $\sigma$ = 1, $\rho$ = 2.5 at an approximate voxel resolution of 20μm. $\sigma$ = 1 suppresses high-frequency noise while preserving gradients at or near the myocyte diameter scale. $\rho$ = 5 corresponds to approximately half the full length of a paediatric myocyte, capturing local orientation without averaging over physiologically distinct regions. Some samples had been previously exposed to fomblin, and as a result, the image quality after the myocyte orientation analysis was sub-optimal. In these samples, $\sigma$ was reduced to 0.25 and therefore essentially removed as the Gaussian filter is truncated at 4 standard deviations and in this case equals 1 pixel, resulting in no averaging effect. Table S2 details structure-tensor parameters used.

Due to the terabyte-scale size of the data, chunk based processing was implemented to enable distributed computation of local tensor fields across a high-performance computing cluster. By splitting computation across multiple processors (parallelization) and optimizing memory management, computation time was reduced from 50 hours to approximately 1-2 hours per 10μm dataset, while maintaining voxel-wise segmentation.

#### Statistical Analysis

To compare ventricular thickness both between ToF and Control specimens and between ventricles, paired t-tests were used. One-way ANOVA was used to assess regional differences in myocyte aggregate orientation between control and ToF groups. In phenotype-specific analysis post-hoc analysis with Tukey’s HSD was performed. HA frequency histograms and transmural profiles were generated for basal, mid-ventricular, and apical levels. One-way ANOVA was also used to compare the shape and mean of HA frequency distributions between phenotypes. To assess the proportion of the right and left ventricular aspects of the interventricular septum, an unpaired t-test was performed.

#### Colourmap selection for Helical Angle Visualization

HA visualization depends critically on the choice of colourmap. The commonly used HSV palette is neither perceptually uniform nor accessible, produces visually dominant yellow bands, and introduces an artificial discontinuity at the –90°/+90° wraparound, particularly impairing interpretation for readers with red–green color-vision deficiency. In line with Society for Cardiovascular Magnetic Resonance recommendations and contemporary scientific visualization standards, we used a cyclic, perceptually uniform HA colormap developed by Herreria (github.com/Pedro-Filipe/cardiac_DTI_colormaps)^7–9^. This palette preserves continuity across angular data, differentiates positive and negative HA, and maintains interpretability across common colour-vision deficiencies, and has been validated in prior diffusion tensor MRI studies (Figure S1).

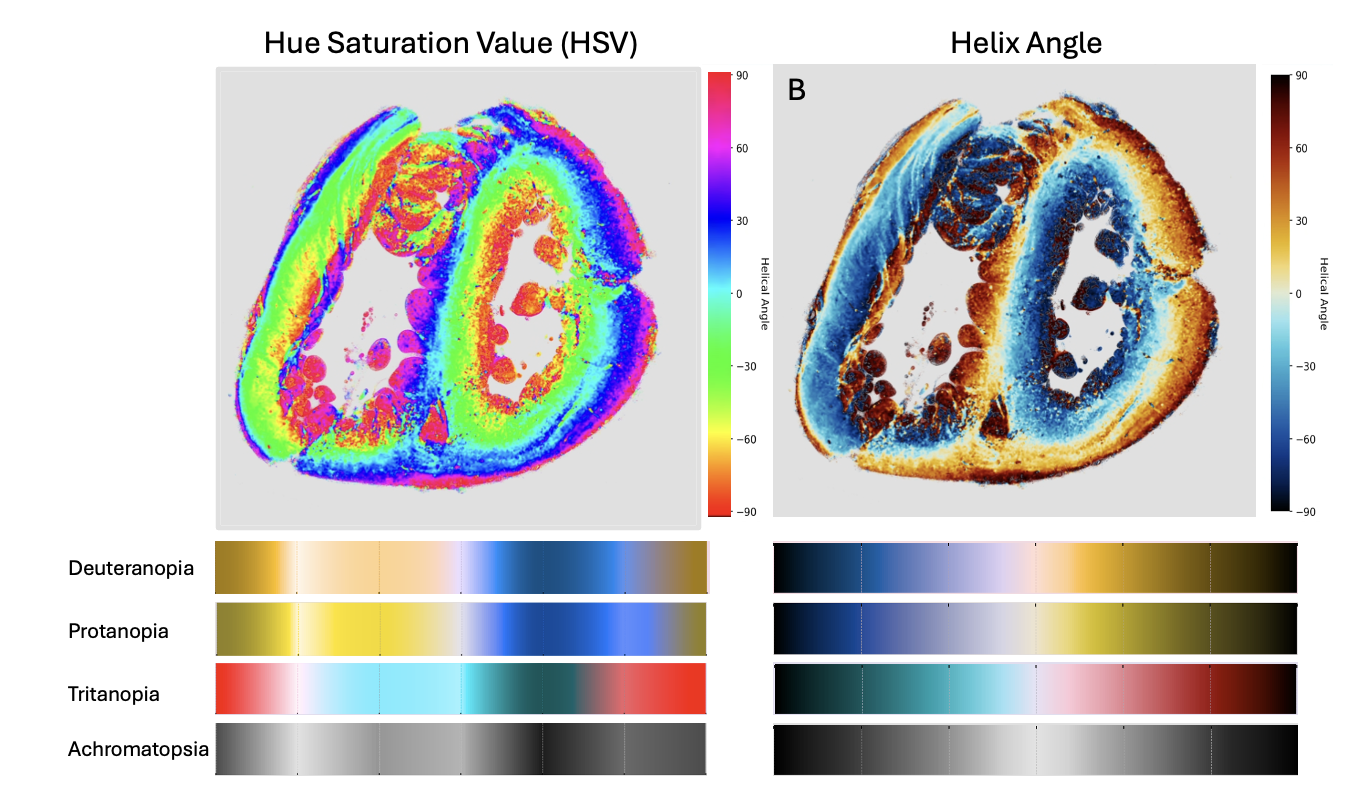

Figure S1: Standard HSV colormap compared to the Helix Angle colourmap used in this study, as viewed by common color vision deficiencies.

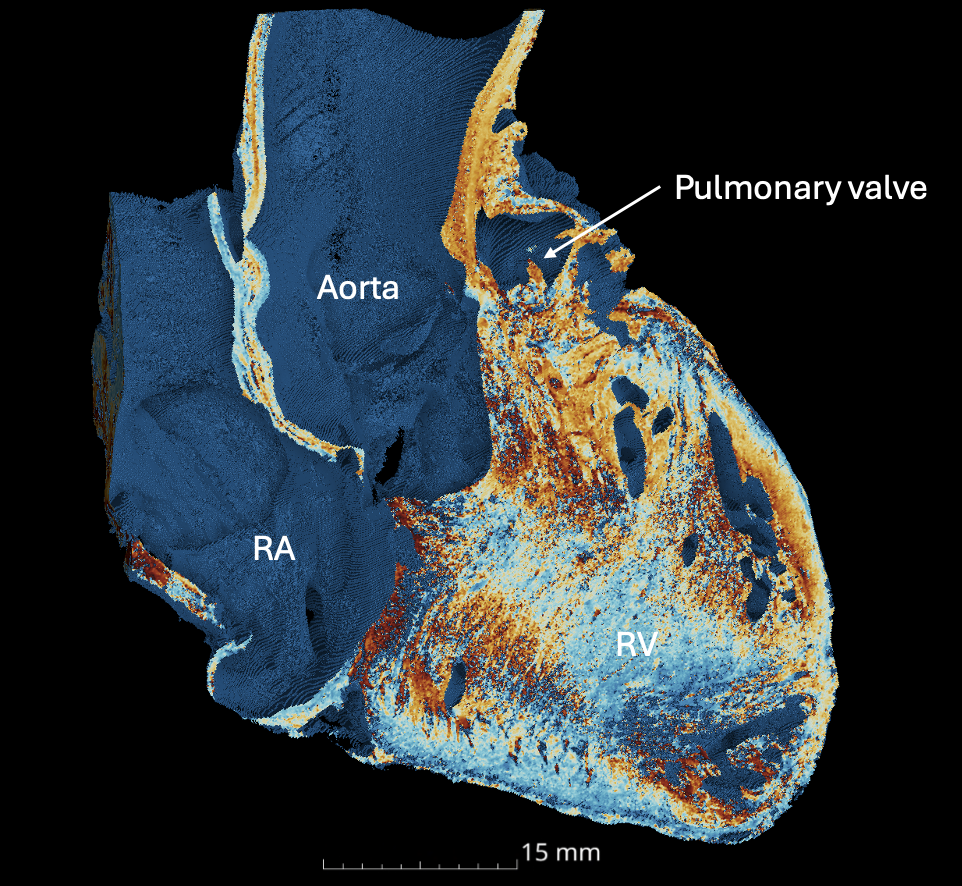

Figure S2: Interventricular septum displayed ‘en-face’ demonstrating the abnormal insertion points extending into the septum from base to apex through the length of the right ventricle (RV). RA – right atrium.

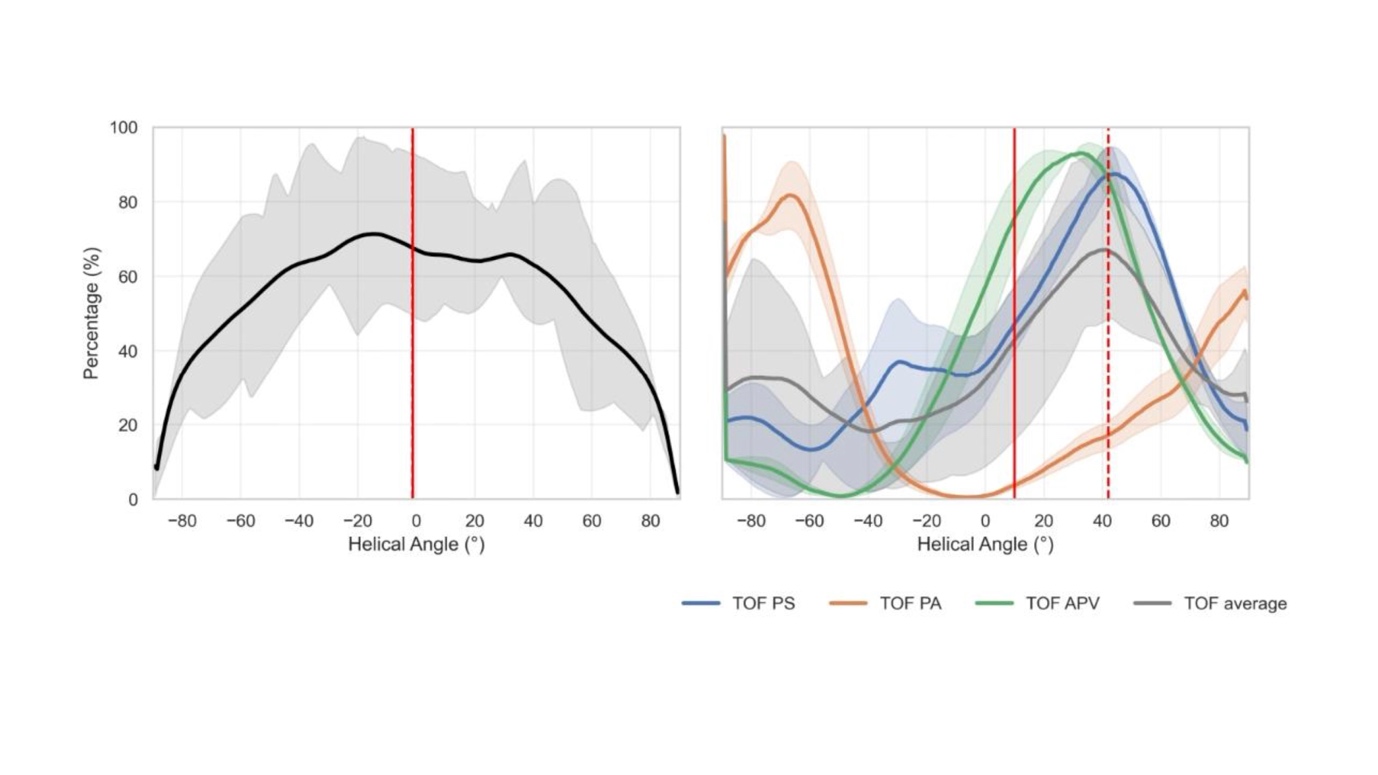

Figure S3: Helical angle frequencies in ToF phenotypes in right ventricular outlet tract across control specimens, classic ToF (pulmonary stenosis), ToF with absent pulmonary valve (APV) and ToF with pulmonary atresia (PA). Interquartile range is shown as the shaded region. Solid red lines mark the numerical means, while circular means are marked by the dashed line.

### Supplementary Tables

Table S1: Scanning parameters for each HiP-CT scan

| Sample ID | DOI | Organ | Voxel size (um) | Propagation distance (m) | Attenuator | Mean energy (keV) | Lateral Field of View (mm) | Projections | Exposure Time (ms) | Accumulation | Acquisition Mode | Scan Time (s/scan) | Total number of scans | Vertical field of view/scan (mm) | Vertical translation |
| --- | --- | --- | --- | --- | --- | --- | --- | --- | --- | --- | --- | --- | --- | --- | --- |
| **125 (T1)** | http://doi.org/10.15151/ESRF-DC-2299323256 | Heart | 9.595 | 10 | Sapphire 5mm Mo 0.21mm | 96 | 81.14 | 12000 | 20 | 1 | Half-acquisition | 265 | 15 | 8.16 | 7 |
| **173 (T2)** | http://doi.org/10.15151/ESRF-DC-2298042950 | Heart | 9.595 | 10 | Sapphire 5mm Mo 0.21mm | 96 | 81.14 | 12000 | 20 | 1 | Half-acquisition | 265 | 14 | 8.16 | 7 |
| **185 (T3)** | http://doi.org/10.15151/ESRF-DC-2298042926 | Heart | 9.595 | 10 | Sapphire 5mm, Mo 0.21mm | 96 | 81.14 | 12000 | 20 | 1 | Half-acquisition | 265 | 15 | 8.16 | 7 |
| **450 (T4)** | http://doi.org/10.15151/ESRF-DC-2298042944 | Heart | 9.595 | 10 | Sapphire 5mm Mo 0.21mm | 96 | 81.14 | 12000 | 20 | 1 | Half-acquisition | 265 | 18 | 8.16 | 7 |
| **727 (T5)** | http://doi.org/10.15151/ESRF-DC-2299337101 | Heart | 9.595 | 10 | Sapphire 5mm, Mo 0.21mm | 96 | 81.14 | 12000 | 20 | 1 | Half-acquisition | 265 | 16 | 8.16 | 7 |
| **769 (T6)** | http://doi.org/10.15151/ESRF-DC-2299323263 | Heart + lungs | 20.01 | 10 | Sapphire 10mm, Ag 0.2mm, C 25mm | 108 | 117.9 | 12000 | 10 | 4 | Quarter-acquisition | 530 | 15 x 2 | 8.00 | 7 |
| **1770 (T7)** | http://doi.org/10.15151/ESRF-DC-2298042938 | Heart | 9.595 | 10 | Sapphire 5mm Mo 0.21mm | 96 | 81.14 | 12000 | 20 | 1 | Half-acquisition | 265 | 15 | 8.16 | 7 |
| **1771 (T8)** | http://doi.org/10.15151/ESRF-DC-2299337089 | Heart | 9.595 | 10 | Sapphire 5mm Mo 0.21mm | 96 | 81.14 | 12000 | 20 | 1 | Half-acquisition | 265 | 15 | 8.16 | 7 |
| **1840 (T9)** | http://doi.org/10.15151/ESRF-DC-2298042932 | Heart +lungs | 19.19 | 12 | Sapphire 5mm Mo 3.75mm SiO2 bars 25mm | 155 | 137 | 12000 | 13 | 3 | Quarter-acquisition | 516 | 21 x 2 | 8.16 | 7 |
| **2010 (T10)** | http://doi.org/10.15151/ESRF-DC-2299479757 | Heart + lungs | 19.19 | 12 | Sapphire 5mm Mo 3.75mm SiO2 bars 25mm | 155 | 137 | 12000 | 13 | 3 | Quarter-acquisition | 516 | 16 x 2 | 8.06 | 7 |
| **2577 (T11)** | http://doi.org/10.15151/ESRF-DC-2299323270 | Heart | 10.005 | 10 | Sapphire 10mm Ag 0.2mm C 20mm | 105 | 90.6 | 12000 | 20 | 1 | Half-acquisition | 265 | 16 | 8.00 | 7 |
| **2359 (C1)** | http://doi.org/10.15151/ESRF-DC-2299337095 | Heart | 9.595 | 10 | Sapphire 5mm Mo 0.21mm | 96 | 81.14 | 12000 | 20 | 1 | Half-acquisition | 265 | 14 | 8.16 | 7 |
| **2337 (C2)** | http://doi.org/10.15151/ESRF-DC-2299323283 | Heart | 10.005 | 10 | Sapphire 10mm Ag 0.2mm C 20mm | 108 | 90.6 | 15000 | 20 | 3 | Half-acquisition | 1026 | 15 | 8.9 | 7 |
| **2625 (C3)** | http://doi.org/10.15151/ESRF-DC-2406973754 | Heart | 7.032 | 7 | Sapphire 10mm, Ag 0.2mm, glassy carbon 30mm | 105 | 66.5 | 15000 | 20 | 3 | Half-acquisition | 198.6 | 11 | 8.02 | 7 |
| **2627 (C4)** | http://doi.org/10.15151/ESRF-DC-2299323277 | Heart | 7.032 | 7 | Sapphire 10mm, Ag 0.2mm, glassy carbon 30mm | 105 | 66.5 | 15000 | 20 | 3 | Half-acquisition | 198.6 | 12 | 8.02 | 7 |
| **2628 (C5)** | http://doi.org/10.15151/ESRF-DC-2299323289 | Heart | 7.032 | 7 | Sapphire 10, Ag 0.3mm, glassy carbon 30mm | 105 | 66.5 | 15000 | 20 | 3 | Half-acquisition | 198.6 | 12 | 8.02 | 7 |
| **2629 (C6)** | http://doi.org/10.15151/ESRF-DC-2299323295 | Heart | 7.032 | 7 | Sapphire 10mm, Ag 0.2mm, glassy carbon 30mm | 105 | 66.5 | 15000 | 20 | 3 | Half-acquisition | 198.6 | 9 | 8.02 | 7 |
| **2630 (C7)** | http://doi.org/10.15151/ESRF-DC-2299323301 | Heart | 7.032 | 7 | Sapphire 10mm, Ag 0.2mm, glassy carbon 30mm | 105 | 66.5 | 15000 | 20 | 3 | Half-acquisition | 198.6 | 9 | 8.02 | 7 |
| **3340 (C8)** | http://doi.org/10.15151/ESRF-DC-2299337064 | Heart | 10.045 | 10 | Sapphire 7mm, Ag 0.3mm | 100 | 90.6 | 15000 | 15 | 1 | Half-acquisition | 248 | 1 | 10.28 | 5 |
| **3341 (C9)** | http://doi.org/10.15151/ESRF-DC-2299337057 | Heart | 7.013 | 7 | Sapphire 7mm, Ag 0.2mm | 92.3 | 57.9 | 14000 | 18 | 1 | Half-acquisition | 277 | 1 | 11.2 | 6 |

Table S2: Rho and Sigma parameters used in structure-tensor calculation.

| Sample | Voxel Size (μm) | $\sigma$ | $\rho$ |
| --- | --- | --- | --- |
| T1 | 9.595 | 1 | 5 |
| T2 | 9.595 | 1 | 5 |
| T3 | 9.595 | 1 | 5 |
| T4 | 9.595 | 1 | 5 |
| T5 | 9.595 | 1 | 5 |
| T6 | 20.01 | 1 | 2.5 |
| T7 | 9.595 | 1 | 5 |
| T8 | 9.595 | 1 | 5 |
| T9 | 19.19 | 1 | 2.5 |
| T10 | 19.19 | 1 | 2.5 |
| T11 | 10.01 | 1 | 5 |
| C1 | 9.595 | 1 | 2.5 |
| C2 | 10.01 | 1 | 5 |
| C3 | 7.032 | 0.25 | 5 |
| C4 | 7.032 | 0.25 | 5 |
| C5 | 7.032 | 0.25 | 5 |
| C6 | 7.032 | 0.25 | 5 |
| C7 | 7.032 | 0.25 | 5 |
| C8 | 10.05 | 0.25 | 5 |
| C9 | 7.02 | 0.25 | 5 |

Table S3: Contrast-to-noise ratios and image quality for specimens associated with prior exposure to fomblin.

| ID | Mean CNR ± SD | Prior exposure to fomblin |
| --- | --- | --- |
| T1 | 11.94 ± 1.74 | No |
| T2 | 11.18 ± 0.78 | No |
| T3 | 11.12 ± 0.89 | No |
| T4 | 11.38 ± 0.59 | No |
| T5 | 3.59 ± 0.25 | No |
| T6 | 10.86 ± 0.31 | No |
| T7 | 15.75 ± 1.50 | No |
| T8 | 5.89 ± 0.49 | No |
| T9 | 12.59 ± 1.11 | No |
| T10 | 5.22 ± 0.88 | No |
| T11 | 12.42 ± 0.94 | No |
| C1 | 10.64 ± 0.58 | No |
| C2 | 2.57 ± 0.03 | Yes |
| C3 | 4.66 ± 1.69 | Yes |
| C4 | 6.89 ± 1.27 | Yes |
| C5 | 6.12 ± 0.37 | Yes |
| C6 | 5.87 ± 0.19 | Yes |
| C7 | 3.89 ± 0.69 | Yes |
| C8 | 8.19 ± 0.69 | No |
| C9 | 7.32 ± 0.32 | No |
